# What one sees depends on how far the eye has moved

**DOI:** 10.1101/2025.09.17.676280

**Authors:** Y. Howard Li, Michele A. Cox, Jonathan D. Victor, Michele Rucci

## Abstract

Humans explore visual scenes through frequent, rapid gaze shifts known as saccades. These movements redirect the high-acuity region of the retina toward objects of interest, thus selecting information based on location. Here, we show that saccade amplitude provides a separate and complementary form of selection, effectively filtering visual information by spatial frequency rather than location. Specifically, a reduction in saccade amplitude attenuates post-saccadic visual sensitivity in an amplitude-dependent range of low spatial frequencies. This effect is highly robust, so that even minute changes in saccade size considerably affect visibility. We show that this phenomenon arises from the way the magnitude-dependent kinematic characteristics of saccades transform the visual world into a spatiotemporal flow: post-saccadic visibility closely follows theoretical predictions based on the spatial information that saccade transients convey within the temporal bandwidth of retinal sensitivity. Thus, saccades not only guide selection based on location, but also filter visual information based on content, actively shaping perception.

## Introduction

Sensory systems tend to be highly sensitive to changes in their input signals and often leverage these variations to establish sensory representations. This strategy makes sense, as input fluctuations hold significant evolutionary value, providing important information about the motion of objects in the scene, such as the movement of a prey or a predator. Moreover, as an organism moves, even stationary objects generate dynamic sensory signals, offering valuable cues about their location and shape. It is, thus, not surprising that use of spatial information derived from self-motion has been widely documented across various sensory modalities, including audition^1,2^, somatosensation^3,4^, and olfaction^5^.

In the visual modality, where two-dimensional space is already explicit on the retina, extraction of spatial information from the temporal changes due to eye or head movement, has been primarily investigated in the context of depth perception, where it provides a powerful cue^6–8^. However, it has long been argued that this encoding strategy may play a more fundamental role in the visual system^9–14^. This is because the visual system responds strongly to temporal changes^15–19^, and humans incessantly move their eyes, even in the periods in between saccades (Fig. 1a-c), when a slow meandering eye motion, known as ocular drift, continually modulates the visual signals experienced by photoreceptors ^20–22^. By transforming a stationary visual scene into a spatio-temporal luminance flow on the retina, ocular drift delivers spatial information in the temporal frequency range of highest neuronal sensitivity^23^. Considerable evidence supports the hypothesis that the visual system uses these input fluctuations to encode space, including impairments in pattern discrimination in the absence of fixational motion^24–26^ and a tight coupling between the strength of fixational modulations and visual sensitivity^27,28^.

**Figure 1:**
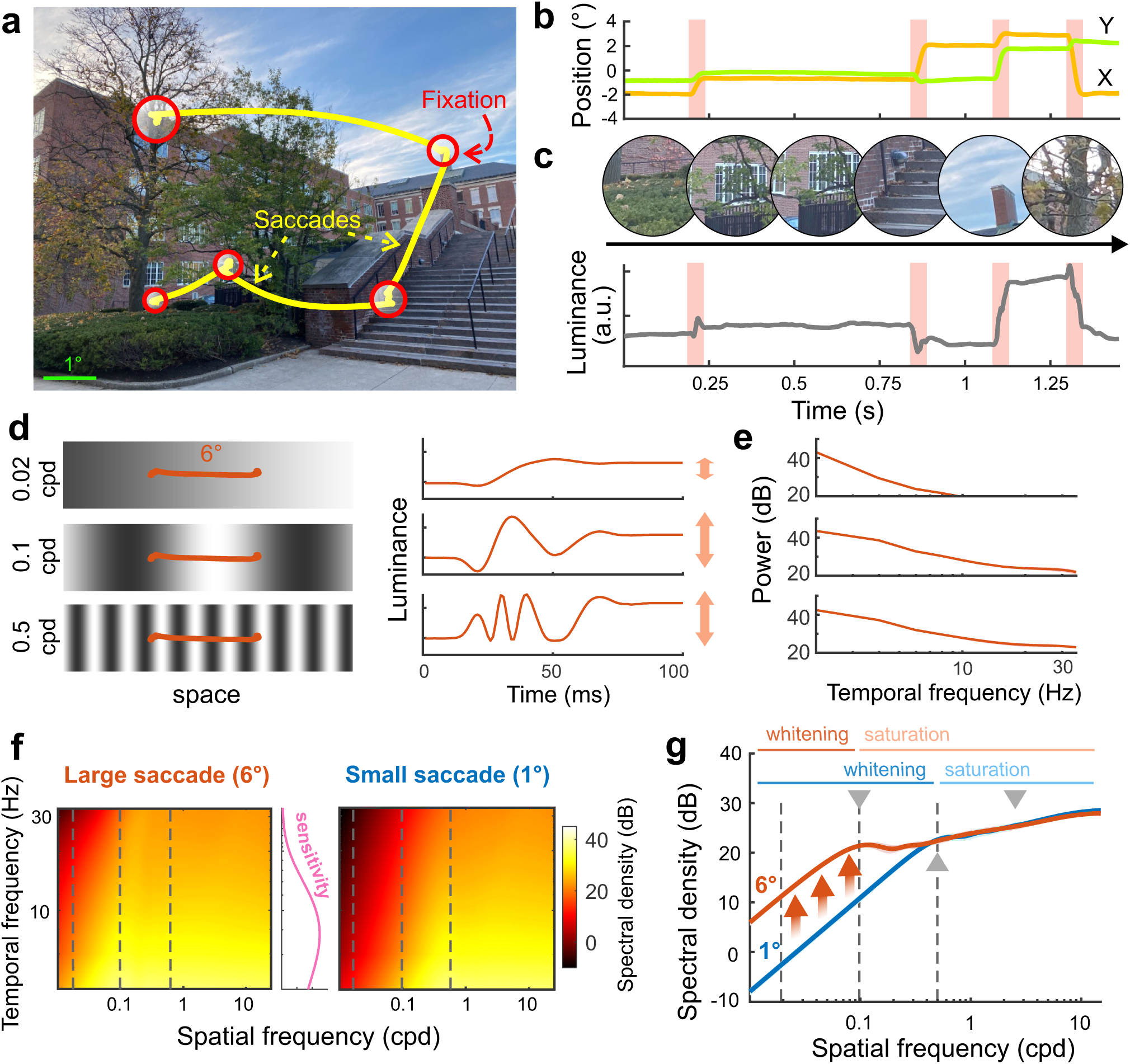
Visual input signals resulting from saccades. **(a)** An example eye trace superimposed on the observed scene. During natural viewing, saccades (yellow lines) continually alternate with periods of fixation (red circles). **(b)** The same trace shown as a function of time. Both horizontal (green line) and vertical gaze displacements (orange) are shown. Pink regions mark the periods of saccades. **(c)** Eye movements modulate input luminance signals. The visual flow experienced by a circular patch on the retina (1*^◦^* diameter; top panel) as the eye moves following the trace in *b*. The bottom trace shows the average luminance. Note the abrupt changes introduced by saccades. **(d)** The luminance transients delivered by saccades depend on the spatial frequency of the stimulus. A 6*^◦^* amplitude saccade (red trace) over gratings of three spatial frequencies (left panel) and the resulting luminance modulations (right panel). **(e)** Power spectra of the luminance transients in *d* averaged over many saccades and starting positions. **(f)** Power spectra of the luminance transients resulting from 6*^◦^* saccades (left panel) and 1*^◦^* saccades (right panel) as a function of the spatial frequency of the stimulus (abscissa). The three dashed lines represent the distributions in *e*. **(g)** Net resulting power within the range of human temporal sensitivity (the pink curve in *f* ^43^). The spectral distributions in *f* are here weighted by temporal sensitivity and integrated across temporal frequencies. Note the presence of a frequency band in which power increases with spatial frequency before saturating (whitening region). The extent of this region depends on saccade amplitude and is wider for smaller saccades. The gray triangles (upward-pointing: Reference; downward-pointing: Probe) mark the spatial frequencies of the stimuli used in this study.

Although research in this area has primarily focused on the input modulations caused by ocular drifts, several observations suggest that a similar strategy of active space-time encoding applies to the signals resulting from other types of eye movements as well. During normal examination of a visual scene, the most prominent luminance transients arise not from fixational movements, but from saccades^29–33^, which—with their well-defined kinematics^34,35^—yield highly structured signals to the retina. These transients differ from the modulations caused by ocular drifts: since saccades induce larger and faster shifts in the retinal image, they cause pronounced input changes for stimuli at low spatial frequencies^36^, where drifts have little effect. In keeping with this observation, saccades have been found to enhance low-frequency sensitivity relative to equivalent periods of eye drifts^37^. Furthermore, coarse-to-fine processing dynamics ^38–41^ have been observed during the typical saccade-drift cycle^42^, as one would expect if the visual system integrates the spatial information conveyed by the temporal signals resulting first from saccades and then from drifts.

The proposal that the visual system extracts spatial information from saccade transients carries important consequences. The way saccades reformat spatial patterns into temporal signals depends primarily on the saccade kinematics, which varies systematically with the amplitude of the saccade itself (Fig. 1c-g). This observation, explained in detail below, leads to the new hypothesis that what one sees following a saccade critically depends on how far the eyes have moved. Specifically, as the saccade size increases, visibility is expected to improve within an *amplitude-dependent* band of low spatial frequencies. In this study, we directly test this hypothesis by measuring the visual performance of human observers and examining whether the signals delivered by saccade transients within the temporal bandwidth of visual sensitivity predict post-saccadic visibility.

## Results

Figure 1 shows how saccades transform a spatial scene into a spatiotemporal flow on the retina. We first focus on the input signals delivered by a saccade of a given amplitude, here a 6*^◦^* saccade (the red trace in Fig. 1d). As illustrated in Fig. 1d, the luminance modulation resulting from the saccade depends on the spatial frequency of the stimulus. At low spatial frequencies, as the eye moves from one position to another, the saccade yields little change (top panel in Fig. 1d). As the frequency increases, the visual transient also increases both in amplitude and speed (middle panel). At sufficiently high spatial frequencies, once the saccade covers more than one cycle of the stimulus, the amplitude of the modulation saturates, but the rate of change continues to increase (bottom panel), spreading power to progressively higher temporal frequencies (Fig. 1e).

These effects are demonstrated more comprehensively in the left panel of Fig. 1f, which shows the power spectrum resulting from a 6*^◦^* saccade as the spatial frequency of the stimulus varies systematically. Only part of the modulation resulting from the saccade falls within the temporal range of visual sensitivity (middle panel in Fig. 1f), as a portion of the signal occurs at temporal frequencies that are too high for the visual system to follow. Fig. 1g provides an estimate of the strength of the input modulation that is effective in driving visual responses, estimated as the total integrated power weighted by the human temporal sensitivity function ^43^(Fig. 1g, orange curve). Note that two distinct regions are present: in a low-frequency band, power increases proportionally to the square of spatial frequency. We will refer to this band as the “whitening” region, as the resulting luminance modulations in this frequency range tend to counterbalance the power spectrum of natural scenes. Beyond a critical frequency, power saturates and no longer depends on the spatial frequency of the stimulus (the “saturation” region).

Importantly, the specific characteristics of the redistribution of stimulus power vary with the amplitude of the saccade. A well-known kinematic feature of saccades is the tight relation between amplitude and speed, the so-called main sequence characteristic^34^. Since larger saccades are also faster, their overall durations vary little with amplitude, and the traces of saccades of distinct sizes differ primarily by a spatial—but not a temporal—scaling factor^36^. Because of the inverse relation between scaling in space and spatial frequency (a spatial expansion corresponds to a shrinkage in frequency), the width of the whitening region follows an inverse proportionality relation with saccade amplitude. Thus, a small saccade, like the 1*^◦^* saccade shown in the right panel of Fig. 1f, exhibits a wider whitening region than the previous 6*^◦^* saccade (Fig. 1g, blue curve), reaching saturation at a larger cut-off frequency and delivering less power at every spatial frequency within the whitening band. Thus, since the extent of the whitening region depends on saccade amplitude, there is a range of spatial frequencies in which larger saccades deliver stronger luminance modulations, as marked by the red arrow in Fig. 1g.

This interaction between saccade amplitude, spatial frequency, and the resulting temporal modulation provides a direct way to examine the influence of saccade transients on spatial sensitivity. Two predictions arise directly from the spectral distributions shown in Figs. 1f, g. First, larger saccades should enhance the visibility of low-spatial-frequency stimuli compared to smaller saccades. Second, smaller saccades should facilitate the perception of stimuli in their saturation band by reducing power at lower frequencies relative to larger saccades. We test these predictions in two separate experiments.

In a forced-choice procedure, subjects were asked to report which one of two orthogonal gratings was more visible following saccades of either 1*^◦^* or 6*^◦^* amplitude (Fig. 2a). The two gratings were superimposed onto each other. One of them—the “Reference”— was always presented at a fixed contrast and frequency (0.5 cycles per degree; cpd). The other—the “Probe”—varied in both spatial frequency and contrast: it was either at lower (0.1 cpd; Low-Probe condition) or higher (2.5 cpd; High-Probe condition) frequency than the Reference (Fig. 2b), and its contrast varied randomly across trials. We estimated the visibility of the Probe relative to the Reference as a function of the Probe contrast.

**Figure 2:**
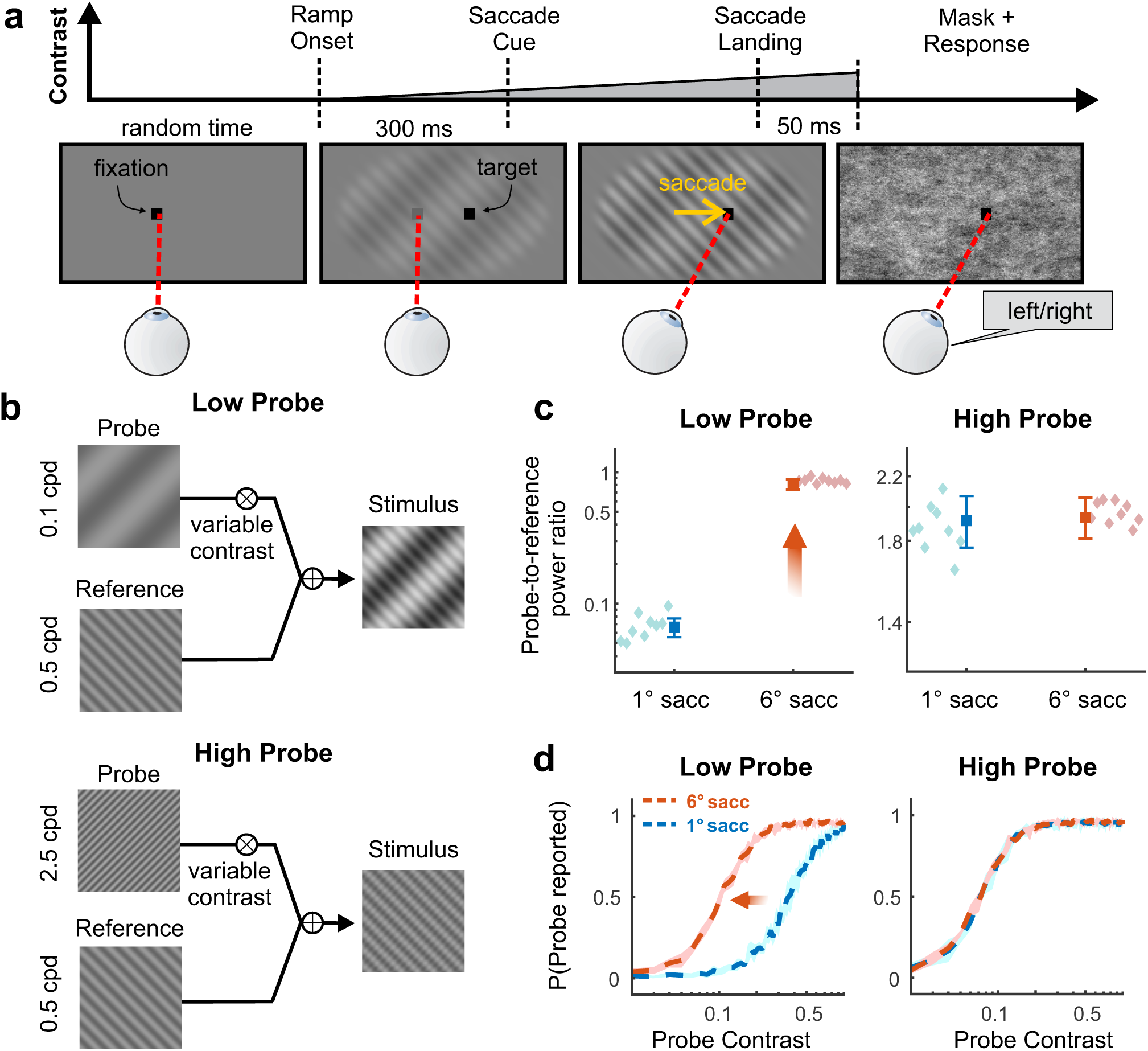
Experimental design and theoretical predictions. **(a)** Subjects made an instructed saccade (amplitude 1*^◦^* or 6*^◦^*) over a stimulus composed of two superimposed orthogonal gratings. The stimulus slowly ramped up in contrast, starting at the beginning of the trial, and was masked 50 ms after the end of the saccade by a Brown noise field (spectral density ∝ *f ^−^*^2^; *f* spatial frequency). Subjects reported which of the two gratings was more visible. **(b)** One of the gratings was always presented at a fixed contrast (the Reference). The contrast of the other grating (the Probe) varied across trials. The spatial frequency of the Probe was either lower (0.1 cpd; Low-Probe condition) or higher (2.5 cpd; High-Probe condition) than the frequency of the Reference (0.5 cpd). **(c)** Relative strengths of the luminance signals resulting from saccades at the spatial frequencies of the Probe and the Reference. Data points represent the ratio in input power delivered by the Probe and the Reference for both small (1*^◦^*) and large saccade (6*^◦^*). In the Low-Probe condition, larger saccades emphasize the Probe relative to the Reference. This effect does not occur in the High-Probe condition. Diamonds are individual data from the experiment in *A* (N = 9). Squares represent the mean power ratio, averaged across subjects (*N* = 18), given by saccades with similar amplitudes during free-viewing of natural scenes. Error bars are ± one standard deviation. **(d)** Sensitivity predicted by the simulated responses of transient neuronal populations tuned to the stimulus frequencies and orientations. In the Low-Probe condition, the stronger signal delivered by the larger saccade is expected to enhance the visibility of the Probe (red arrow). In contrast, no difference in visibility is expected following 1*^◦^* or 6*^◦^* saccades in the High-Probe condition.

The frequencies of the gratings and saccade amplitudes were chosen so that the luminance transients delivered in the Low-Probe and High-Probe conditions would lead to contrasting predictions. The fixed frequency of the Reference was selected so that it fell in the saturation range for both 1*^◦^* and 6*^◦^* saccades (upper triangle in Fig. 1g). In contrast, the frequency of the Probe was chosen so that the resulting luminance modulations from small and large saccades would differ in one condition, but not in the other. Specifically, in the High-Probe condition, the 2.5 cpd Probe fell within the saturation ranges of both saccades (right-most upside-down triangle in Fig. 1g), thus yielding similar input signals for small and large saccades. In contrast, in the Low-Probe condition, the 0.1 cpd Probe fell within the whitening range of the 1*^◦^* saccade and in the saturation range of the 6*^◦^* saccade (left upside-down triangle in Fig. 1g), thus delivering a stronger input signal for the large saccade. Therefore, whereas the 6*^◦^* saccade always yields identical transients for the Probe and the Reference, with the 1*^◦^* saccade, the Probe gives a weaker transient than the Reference in the Low-Probe condition, but not in the High-Probe condition.

We confirmed that the saccades performed by the subjects participating in our experiments delivered the expected transients by reconstructing the visual flow experienced by their retinas. For each subject, we first estimated the strength of saccade transients by filtering the input signals from the Probe and the Reference by the temporal sensitivity of the human visual system^43^, and then compared the relative impact of the Probe and the Reference by means of their power ratio (the signal-to-noise ratio, SNR; Fig. 1g). As expected, the two saccade amplitudes deliver comparable power ratios with the High-Probe stimulus configuration. In contrast, in the Low-Probe condition, the SNR of a 1*^◦^* saccade is approximately one order of magnitude lower than that of a 6*^◦^* saccade (Fig. 2c). Similar results were obtained by examining the power at the spatial frequencies of interest of the visual transients delivered by saccades of comparable amplitudes collected as subjects freely examined images of natural scenes (bold squares in Fig. 2c).

To provide an intuition of the perceptual consequences of these input signals, we estimated the psychometric functions predicted by a simple ideal-observer model (Fig. 2d). In each trial, the model estimated the average responses of linear filters simulating populations of V1 neurons as they were exposed to reconstructions of the saccade-induced visual transient experienced by the subjects’ retinas. Neurons were tuned to the two orientations and frequencies of the Probe and the Reference, and the model reported as more visible the grating that elicited a higher overall response. Fig. 2d shows the model’s probability of reporting the Probe to be more visible than the Reference as a function of the Probe’s contrast. As shown by these data, small and large saccades led to distinct perceptual consequences in the Low-Probe, but not in the High-Probe, condition. In keeping with the strength of the visual input signals, the large 6*^◦^* saccade improved visibility of the low-frequency probe, as indicated by the leftward shift in the predicted psychometric function (Fig. 2d, left panel). In contrast, with a high spatial frequency Probe, the two saccade amplitudes gave virtually identical psychometric functions, predicting similar visibility (Fig. 2d, right panel).

Perceptual performances measured in our experiments closely followed these theoretical predictions. Fig. 3a-b reports the psychometric functions obtained from one representative subject. In the Low-Probe condition, when the strength of the luminance modulation at the Probe frequency varied with saccade amplitude, the visibility of the Probe was enhanced by a larger saccade, as shown by the leftward shift in the 6*^◦^* psychometric function relative to the 1*^◦^* in Fig. 3a. This effect was highly consistent across observers, with all individual subjects exhibiting a statistically significant improvement in sensitivity (solid lines in Fig. 3C; p *<* 0.03, parametric bootstrapping test). On average across subjects, contrast sensitivity increased by 37% with the larger saccade (Fig. 3c, p *<* 0.001; paired t-test). In contrast, in the High-Probe condition, performance was virtually unaffected by the amplitude of the saccade: the psychometric functions measured with 1*^◦^* and 6*^◦^* saccades largely overlapped with each other (Fig. 3b), resulting in similar Probe sensitivity in the two conditions (Fig. 3d; p = 0.95).

**Figure 3:**
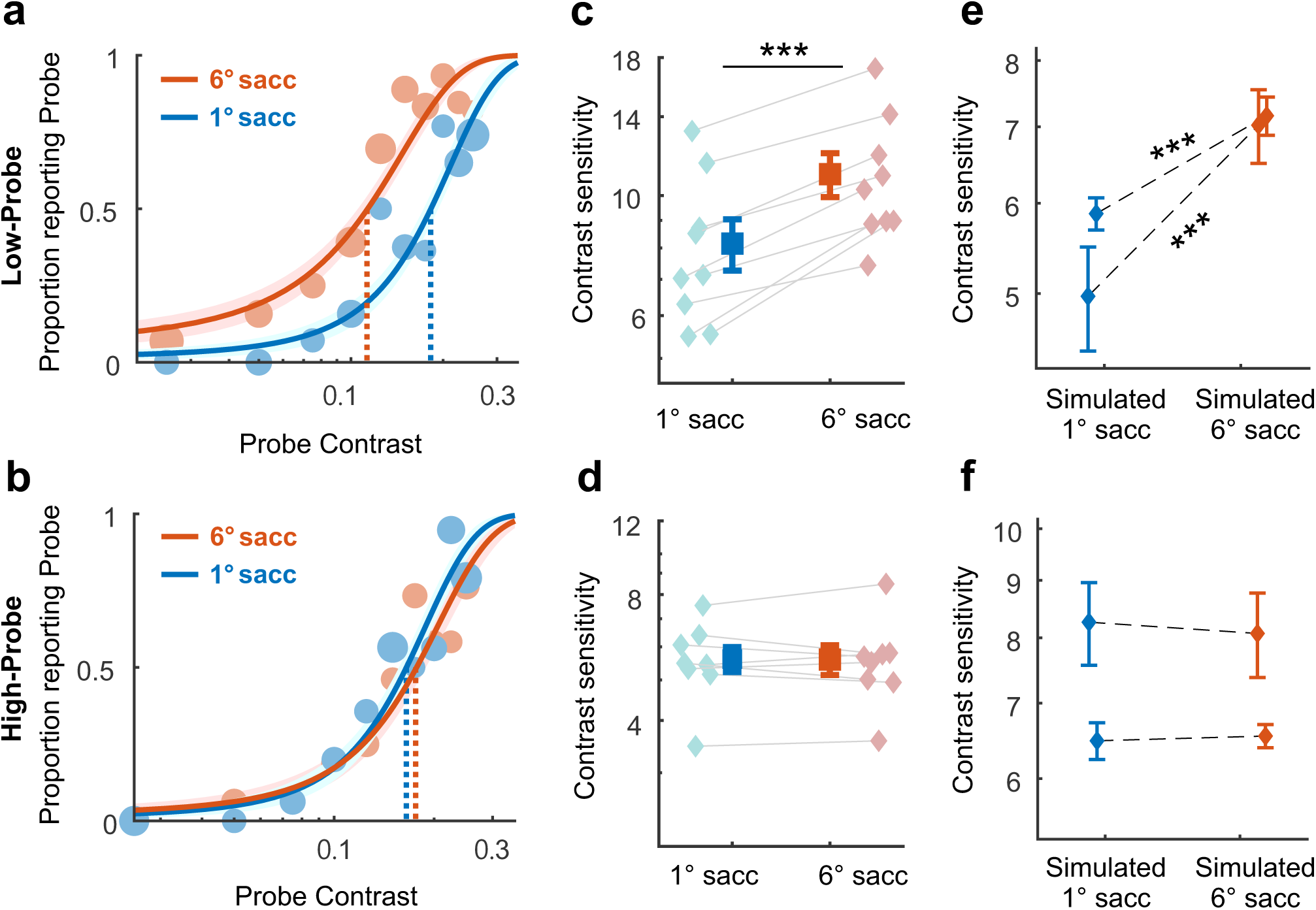
Experimental results. **(a-b)** Measured psychometric functions for an individual subject. The two curves mark the probability of reporting the Probe as a function of its contrast for both small (1*^◦^*, blue) and large (6*^◦^*, red) saccades, and marker size indicates trial numbers at each contrast level. Dashed lines mark the point of subjective equality (the 50% threshold). Shaded areas represent ± one standard error. *(a)* Large saccades enhance probe visibility when the Probe is at a lower spatial frequency than the Reference (red arrow; p *<* 0.001, parametric bootstrap test). *(b)* In contrast, saccade amplitude has no influence when the Probe is at a higher spatial frequency than the Reference (p = 0.69). **(c-d)** Comparing contrast sensitivity for small and large saccades. Bold symbols represent averages across subjects (*N* = 9). A 6*^◦^* saccade enhances visibility in the Low-Probe (*c*, p *<* 0.001, paired t-test) but not in the High-Probe condition (*D*, p = 0.95). Error bars represent standard errors. Diamonds are the data from individual subjects. **(e-f)** Contrast sensitivity with simulated saccades. Data are shown for two subjects. Error bars represent ± one standard error.

It is worth noting that the discrepant perceptual effects reported in the Low- and High-Probe conditions are not due to differences in behavior or extraretinal differences related to saccade production. Specifically, results were not caused by small differences in the reaction times of small and large saccades (approximately 50 ms longer in the 1*^◦^* saccade), as demonstrated in Fig. S1 by considering only pools of trials with comparable reaction times for both saccade amplitudes. Furthermore, similar results were also obtained in a control experiment with passive exposure to saccade motion, a condition that eliminates possible extra-retinal influences (Fig. 3e-f, also see Fig. S2). In this experiment, rather than performing saccades, observers maintained fixation while the stimulus moved on the display following previously recorded saccade traces. Simulated saccades delivered the same retinal motion as real ones, but lacked associated extra-retinal signals, as no motor command was generated. Yet, as with real saccades, the larger simulated saccade selectively improved visibility at low spatial frequencies, confirming that the effects result from the way that the stimuli move on the retina.

The results in Figs. 1-3 show that visibility follows the spatiotemporal power of the input stimulus delivered by saccades. Because of the existence of a whitening region in which saccade transients counterbalance the power spectrum of natural scenes, the effective strength of stimuli in a low spatial frequency band is attenuated on the retina. Critically, as observed in Fig. 1g, the bandwidth of the whitening region depends on the saccade amplitude. This observation leads to our second prediction: a saccade smaller than 1*^◦^* is expected to exert an influence opposite to that of the large saccade in Fig. 3, enhancing visibility of the Probe when it is at higher—rather than lower—frequency of the Reference.

This prediction is explained in detail in Fig. 4a. In the previous experiment, the perceptual enhancement was caused by the greater power delivered by a larger saccade at low spatial frequencies: whereas the Reference and the High-Frequency Probe consistently remained in the saturation region for both saccade amplitudes, the Low-Frequency Probe transitioned from the whitening region of the 1*^◦^* saccade to the saturation region of the 6*^◦^* saccade, resulting in a stronger visual signal. Note, however, that the bandwidth of the whitening region increases as saccade amplitude decreases (Fig. 1g). This means that a sufficiently small saccade (like the 0.4*^◦^* saccade in Fig. 4) would further extend the whitening region to encompass the Reference. This phenomenon should weaken the transient from Reference, affecting the relative strength of the visual input signals from the two components of the stimulus (Fig. 4b) and enhancing the visibility of the Probe (supplementary Fig. S3).

**Figure 4:**
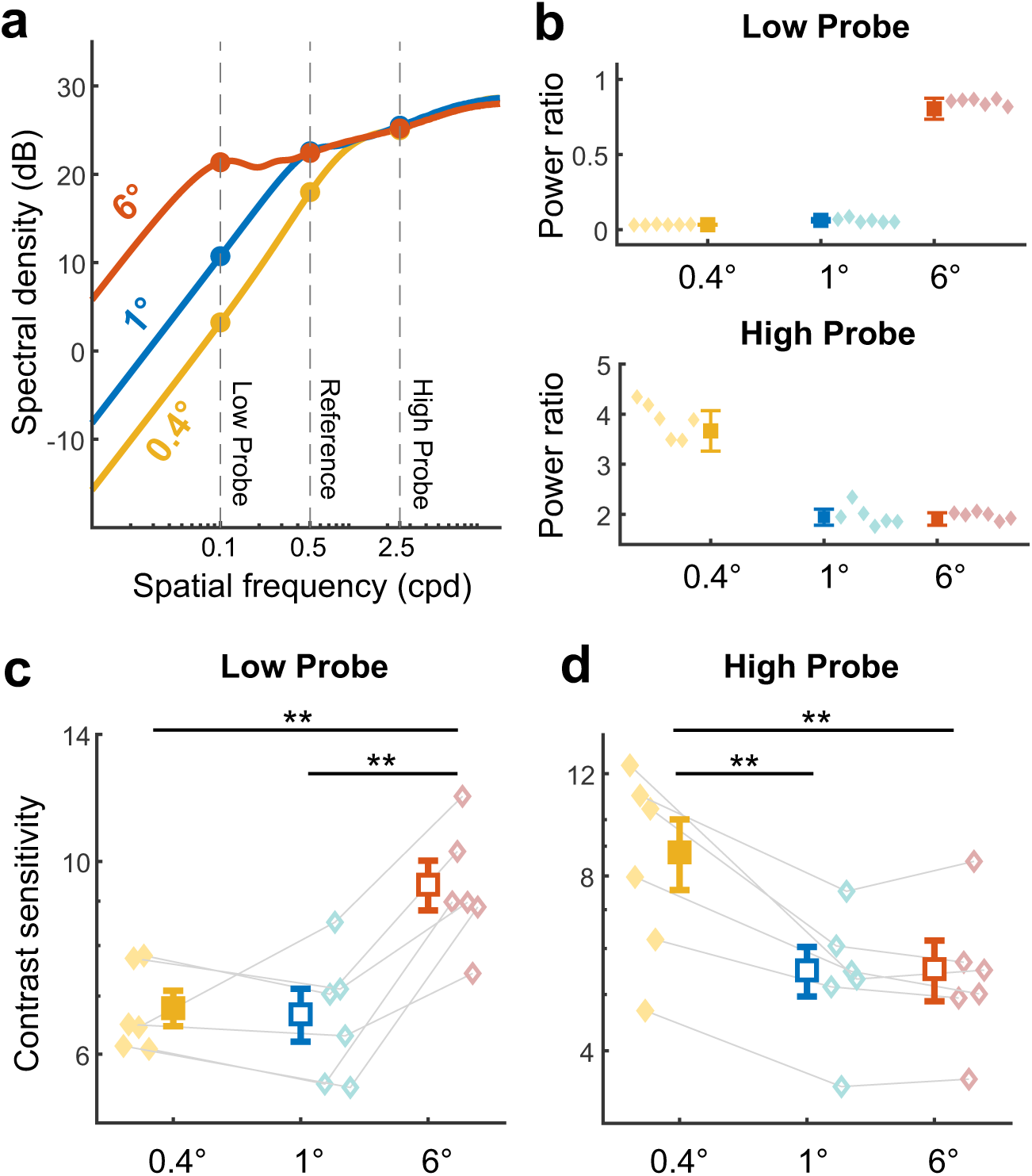
Perceptual consequences of smaller saccades. A saccade smaller than 1*^◦^* is expected to yield a perceptual improvement in the High-Probe condition. **(a)** The stimulus power delivered from the previously tested saccades (1*^◦^* and 6*^◦^*) is here compared to the power resulting from a saccade with 0.4*^◦^* amplitude (yellow; *cf*. Fig.1*g*). The smaller saccade extends the region of difference in power, with reduced power delivered by the Reference (yellow arrow). **(b)** Ratio of power delivered by the Probe and reference. A 0.4*^◦^* saccade enhances the signal resulting from the Probe in the High-Probe condition (yellow arrow), when the Probe is at higher spatial frequency than the Reference, but not in the Low-Probe condition. Graphic conventions are as in Fig. 2c. **(c-d)** Sensitivity measured following saccades with the three amplitudes. In the Low-Probe condition (left), a 6*^◦^* saccade improves probe visibility compared to the 1*^◦^* (p = 0.005) and the 0.4*^◦^* (p = 0.003) saccade. In the High-Probe condition (right), a 0.4*^◦^* saccade selectively improves visibility (p = 0.002 compared to both the 1*^◦^* and 6*^◦^* saccade). Graphic conventions are as in Fig. 3c-d. Open symbols represents data points replotted from Fig. 3c-d for the *N* =6 participants in this experiment.

To test this prediction, we assessed the visibility of the Probe following a small saccade of 0.4*^◦^* amplitude. Fig. 4c compares these new data to the visibility measurements obtained with the larger saccades 1*^◦^* and 6*^◦^*, replotted here from Fig. 3 for the subset of subjects who participated in this second experiment. In the Low-Probe condition, the condition in which a large saccade is beneficial, performance with 0.4*^◦^* and 1*^◦^* saccades were virtually identical. Importantly, however, as predicted by the strength of saccade transients, the 0.4*^◦^* saccade was beneficial in the High-Probe condition, significantly improving the visibility of the Probe when it was at lower frequency than the Reference. This perceptual improvement was highly consistent across subjects: all participants exhibited significantly higher performance with a 0.4*^◦^* saccade than a 1*^◦^* saccade (p *<* 0.002, bootstrap tests). This effect is remarkable given that it occurs with a mere half-degree difference in saccade amplitude. It shows that even small changes in eye movements carry important consequences on visual functions.

## Discussion

Humans explore visual scenes by making saccades of various amplitudes. These eye movements serve the primary function of redirecting the high-acuity fovea towards objects of interest. In doing so, however, saccades also deliver strong luminance transients to the retina. The characteristics of these transients depend on saccade amplitude, since amplitude and saccade kinematics are tightly linked, and the latter determines the spatial information conveyed within the temporal range of sensitivity of retinal neurons by constraining the motion of the stimulus on the retina^37^. In line with this reformatting in visual stimulation, we found considerable and highly consistent changes in spatial sensitivity when observers made saccades of different sizes. Specifically, our results show that increasing saccade amplitude enhances visibility in a low range of spatial frequencies and decreases the bandwidth in which this effect occurs; conversely, progressively smaller saccades increase the relative visibility of higher spatial frequencies by attenuating power at low spatial frequencies. These findings highlight the importance of oculomotor-induced temporal transients in establishing spatial representations and suggest that saccades provide not only spatial selection, but also spatial frequency selection.

Support for a role of eye movements in reformatting spatial information into a useful spatiotemporal flow comes from previous studies that examined visual sensitivity and oculomotor behavior at fixation. During fixation, a persistent eye motion, known as ocular drift, continually shifts the retinal image across many photoreceptors following seemingly random trajectories. The resulting visual input signals depend on both the characteristics of eye drift and the stimulus: during viewing of natural scenes, normal fixational drift equalizes the strength of luminance fluctuations (the power at nonzero frequencies) over a broad range of spatial frequencies^23^, a band much wider than the whitening range of saccades. Previous studies have shown that, in accordance with oculomotor influences on input signals, sensitivity to high spatial frequencies is selectively impaired in the absence of the retinal image motion caused by ocular drift^24,25^. In fact, spatial sensitivity closely follows the strength of the resulting luminance modulations ^28^, and the characteristics of drift motion performed when subjects are just asked to maintain steady gaze on a point reliably predict individual acuity limits^27^. Furthermore, humans tune their eye drifts according to the task in a way that enhances the resulting luminance modulations ^26,44^. By showing that a similar encoding scheme extends to the temporal transients from saccades, the present study suggests a general strategy of active visuomotor encoding: visual representations do not just rely on the image on the retina, but also on how visual signals change because of eye movements, *i.e.*, on the full oculomotor-shaped spatio-temporal flow of visual information.

Our data demonstrate that the amplitude-dependent perceptual modulations induced by saccades are highly robust. All participants in our experiments exhibited improved performance under conditions where input signals from saccades were expected to enhance discrimination (Figs. 3c, e, and 4c, d). Interestingly, while these perceptual effects are substantial, they are attenuated relative to the pronounced impact of saccades on visual input signals. For example, saccades with amplitudes of 1*^◦^* and 6*^◦^* produce transients that differ by nearly an order of magnitude in power (Fig. 2c), which our linear model predicts should result in a threefold perceptual improvement (Fig. 2D). However, empirical sensitivity measurements only improved by approximately 37% (Fig. 3c). This discrepancy may arise from several factors, including attenuation due to saccadic suppression—specifically, its retinal component given the results with simulated saccades in Fig. 3e,f—, normalization mechanisms in cortical responses ^45^, and processes related to perceptual constancy^46^. A similar compression in sensitivity occurs when comparing the contrast of supra-threshold stimuli at different spatial frequencies, where judgments of relative contrast are much closer to veridical than would be suggested by their ratios to threshold ^47,48^. This effect aligns with the visual system’s overarching goal of estimating the physical contrast of a stimulus while discounting variations in transient strength caused by saccades of different amplitudes.

Previous studies have reported various types of visual enhancements immediately following saccades. For instance, both contrast sensitivity and pattern discriminability improve in a short period after saccade landing^49–51^. Ocular-following, the tracking of a suddenly moving object, is also facilitated when the motion starts right after a saccade rather than at fixation^52–54^. Perceptual modulations around saccades have been commonly attributed to extra-retinal processes related to attention ^55–57^, saccade planning^58^ and execution^59,60^.

However, the visual input signals resulting from saccades are also important^61,62^, and the luminance changes introduced by saccades have been reported to play a role for ocular following^54,63^, visual stability^64,65^, and contrast sensitivity^37,42^. For contrast sensitivity, rapid gaze shifts have been shown to be specifically beneficial for visibility of low spatial frequencies, as one would expect from the stronger signals that saccades deliver relative to fixational drifts in this frequency range^42^. Our results go beyond the previous literature by showing that the amplitude of a saccade matters: post-saccadic visibility critically depends on saccade amplitude, as predicted by the way saccades of different size redistribute spatial information in the temporal domain, *i.e.*, the strength of the input modulations delivered within the temporal bandwidth of retinal neurons. It remains to be determined how and where these oculomotor-shaped luminance transients are combined with the associated extra-retinal signals to establish spatial representations^66^.

The post-saccadic enhancements shown in our experiments should not be confused with the modulations responsible for saccadic omission and suppression: the lack of awareness to the inter-saccadic motion of the retinal image^67–69^, and the reduced sensitivity to stimuli briefly exposed during saccades^50,70,71^. Both phenomena are known to rely not just on extra-retinal signals^72,73^ but also on the visual input to the retina^74–76^. Our finding that sensitivity varies with the characteristics of saccade transients adds new considerations to this interplay. A lingering question regarding saccadic omission is how a relatively weak extraretinal modulation^68,77,78^ could suppress visibility of low spatial frequencies, which is known to increase when the stimulus translates at high speeds^79^. This latter enhancement is consistent with the idea of encoding spatial information in the temporal domain, as a constant-speed translation simply shifts power to a temporal frequency that depends on the spatial frequency of the stimulus and the speed of translation. However, the way saccades move stimuli on the retina differs drastically from uniform motion, and the power spectrum analysis in Fig. 1f –*g* shows that saccades actually yields weak signals at low spatial frequencies. Thus, even a moderate suppression in sensitivity may, by itself, be sufficient to prevent visibility of retinal image motion during saccades.

Regarding saccadic suppression, examination of retinal influences has been traditionally framed in the context of masking, namely how the post-saccadic image perceptually covers the image of the inter-saccadic probe. However, rather than the interaction between the two images, our results emphasizes the need to examine the interference between the luminances transients from the probe and those resulting from eye movements. These transients differ in fundamental ways, as—unlike oculomotor modulations—the brief probe flash preserves the spatial frequency distribution of the probe image at every temporal frequency. This idea of an interference between transients accounts for several reported properties of saccadic suppression. It predicts stronger suppression with low spatial frequency probes^50^ because this is the range where saccades deliver stronger luminance transients than fixational drifts. It also predicts that suppression increases with saccade amplitude^80^, because larger saccades deliver more power at low frequencies, as shown by Fig. 1g and, perceptually, by our results. Thus, consideration of temporal transients may help explaining not only post-saccade enhancements in visibility, but also the reduced sensitivity during saccades, and raises the hypothesis that the extraretinal modulations that are known to contribute to these effects may work synergistically with saccade transients.

The finding that what one sees following a saccade depends on the amplitude of the gaze shift leads to new hypotheses. An emerging possibility is that saccade control may not just be based on spatial selection—the positioning of the high-acuity fovea on locations of interest—, but also, by virtue of the amplitude-dependent strength of saccade transients with stimuli at different spatial frequencies, on a process of selection in spatial frequency (Fig. 1g). In this view, one would expect saccade amplitude to be modulated by the spatial frequency range relevant to the task. This spatial filtering could be used to enhance sensitivity to the spatial frequencies in a target, or to suppress sensitivities to the spatial frequencies in the background or in distractors. Although this hypothesis has not been directly tested, it is consistent with the observation that saccade amplitude increases when searching for low spatial frequency targets in natural scenes^81^. This action enhances saccade transients from the target stimuli. Furthermore, saccade amplitude decreases when a high-pass filter is applied to the scene, which is consistent with the reduced low spatial frequency content^82^.

Our results also emphasize the need to study visual impairments in conjunction with eye movements^83,84^, as inaccurate control of saccade amplitude and/or changes in saccade kinematics that affect their transients are expected to carry perceptual consequences. Abnormal saccades have been reported in multiple conditions that are also known to present visual impairments. For example, the faster and smaller saccades observed in schizophrenia patients^85–88^ are expected to alter the normal coarse-to-fine perceptual dynamics resulting from the saccade-fixation cycle^42^, possibly contributing to the reported visual impairments in figure-ground segmentation and visual acuity ^89,90^. Similarly, the slower saccades observed in some patients affected by autism^91^ are expected to further weaken transients in the whitening frequency band, an effect that could contribute to the reported attenuation in sensitivity at low spatial frequencies ^92^. Further work is needed to investigate these emerging hypotheses.

## Methods

### Subjects

Data were collected from a total of 9 subjects (5 males and 4 females, age range 19-36). All subjects had at least 20/20 uncorrected visual acuity as determined by a standard Snellen eye chart. With the exception of one of the authors, all participants were näıve about the purpose of the experiments and were compensated for their participation. Informed consent was obtained from all subjects as per the approval of the Research Subjects Review Board at the University of Rochester.

### Stimuli and apparatus

Stimuli consisted of two superimposed, orthogonal sine-wave gratings tilted by ±45*^◦^* relative to the horizontal meridian. Stimuli were presented over a uniform gray background (9.1 *cd/m*^2^) and were fully visible through an elliptical window (15*^◦^* by 8*^◦^*) surrounded by region of decreasing transparency (Gaussian envelope with *σ* = 2*^◦^*). One of the gratings, the “Reference”, had a fixed spatial frequency of 0.5 cpd and a fixed Michelson contrast of 0.1. The other grating, the “Probe”, was at either lower (0.1 cpd, Low-Probe condition) or higher frequency than the Reference (2.5 cpd, High-Probe condition), and its contrast varied randomly across trials (0.01 - 0.4 range, method of constant stimuli). Low- and High-Probe trials were intermixed. Stimuli were displayed on a LCD monitor (ASUS258) at a resolution of 1,920 × 1,080 pixels and at a refresh rate of 200 Hz. They were rendered by means of EyeRIS^93^, a custom system for gaze-contingent display control designed to guarantee precise synchronization between the stimulus on the monitor and oculomotor recordings. Stimuli were viewed binocularly.

The movements of the right eye were continuously recorded by means of a digital Dual Purkinje Image eye-tracker (the dDPI ^94^), a system capable of resolving movements smaller than 1*^′^*. Data were acquired in digital form at 1 kHz. To ensure precise measurement of eye movements, the heads of the subjects were immobilized with a custom dental-imprint bite-bar and a head-rest. The monitor was located at a distance of 83 cm from the observer, resulting in each pixel subtending an angle of approximately 1.2*^′^*.

### Procedure

Subjects participated in multiple experimental sessions, each lasting approximately one hour. Data were collected in blocks of 100 trials, each lasting around 10 minutes, with rest periods in between. In every block, subjects performed saccades of a different amplitude. Blocks of trials either alternated between two saccade amplitudes (1*^◦^* and 6*^◦^*; *N* =6 subjects) or three amplitudes (0.4*^◦^*, 1*^◦^* and 6*^◦^*; *N* =6 subjects, 3 new).

Before each block of trials, a gaze-contingent calibration procedure ensured accurate localization of the line of sight. This procedure, described at length in previous publications ^95^ consisted of two phases, with a gaze-contingent refinement following a standard oculomotor calibration. In this second phase, observers fixated again on the nine points of a 3 × 3 grid and corrected possible offsets to the estimated gaze location, which was displayed in real-time on the display. These corrections were then incorporated into a bilinear transformation that mapped oculomotor measurements into screen coordinates.

A trial began with the subject fixating on a marker (a 10*^′^* dot). Once the subject maintained fixation within 1*^◦^* from the marker for longer than 1 s, the stimulus started ramping up in contrast, reaching its maximum in 1 s. A shift in the fixation marker acted as saccade cue, instructing the subject to perform a saccade of a specific amplitude. The saccade cue occurred 300 ms after ramp onset, an interval necessary to ensure that the saccade would deliver a sufficiently strong transient to the retina. The eye velocity was continually monitored in real-time by EyeRIS, and saccade landing was marked as the instant in which the eye speed dropped below 3*^◦^*/s. The stimulus remained visible for 50 ms after saccade landing before being masked by a Brown noise field that covered the entire display (spectral density ∝ *k^−^*^2^, *k*: spatial frequency). The subject reported which of the two orthogonal gratings was more visible by pressing one of two buttons on a joypad.

The experiment with simulated saccades (Figs. 3E-F and S2) followed the same procedure, except that the fixation marker did not move, and subjects maintained fixation at the center of the monitor throughout the trial. After 450 ms waiting time, which accounted for the pre-saccadic exposure in the real-saccade experiment, the gratings moved on the monitor following the horizontal trace of a previously recorded saccade from the same observer, so to replicate the visual input signals delivered by the saccade while eliminating associated extraretinal influences.

### Data Analysis

The recorded oculomotor traces were analyzed offline and segmented based on eye velocity. Eye movements with a speed greater than 3*^◦^*/s and amplitude larger than 10*^′^* were classified as saccades. Only trials with high-quality oculomotor measurements were selected for further analysis. Trials in which the subjects blinked, or made other saccades in addition to the instructed one, or delayed execution of the saccade by more than 700 ms were excluded from data analysis. Furthermore, trials in which subjects did not move the line of sight by the correct amount, either because the landing error was larger than 1*^◦^* or because the saccade amplitude deviated by the requested gaze shift by more than 50%, were also discarded.

For every subject, psychometric functions were estimated by fitting experimental data with cumulative Gaussian functions via a maximum-likelihood procedure^96^. We use the psychometric functions to estimate the 50% performance threshold, the contrast at which the Probe and the Reference were equally visible (the Point of Subjective Equality, PSE). Statistical significance of difference in thresholds between psychometric functions was assessed for each subject via one-sided parametric bootstrapping of the threshold value with 500 repetitions^97^. Sensitivity was measured by taking the inverse of the Probe contrast at the PSE. A within-subject paired *t*-test was used to estimate the statistical significance of the difference of sensitivities as a function of saccade size.

To exclude possible confounds from pre-saccadic exposure to the stimulus ramp, we also examined performance after equalizing the reaction times to small and large saccades (Fig. S1). To this end, for each subject in Fig. 3c-d, we discarded 1*^◦^* saccade trials with long delays and 6*^◦^* saccade trials with short reaction times so to match the mean duration of pre-saccadic exposure. The selected trials were then analyzed as described above.

### Power spectrum analysis and sensitivity prediction

Spectral analyses were conducted following the factorization approach previously described in the literature ^36^. The key quantity is the power redistribution function *Q*(**k***, ω*) resulting from eye movements—in this case a saccade of a given amplitude. *Q* is a function of spatial frequency **k** and temporal frequency *ω*. It determines the temporal power distribution on the retina delivered by movement during viewing of an external stimulus of unit power at spatial frequency **k** (Fig. 1f). Under the plausible assumption of statistical independence between the autocorrelation of the image on the retina and the second-order statistics of gaze displacement, the redistribution function *Q* enables estimation of the spectral density of the retinal input signals, *S*(*k, ω*) resulting from the considered type of eye movements during viewing of arbitrary scenes:

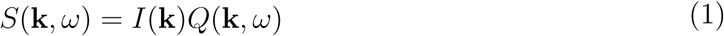

where *I*(**k**) is the power spectrum of the external scene. By separating retinal image statistics and eye movements, this method enables spectral estimation with high temporal resolution at any desired spatial frequency **k**.

For every subject and saccade amplitude, *Q*(**k***, ω*) was estimated based on 100 recorded saccade traces. Each saccade trajectory, ***ξ***(*t*), was isolated in a 512 ms segment that only contained the considered movement. Eye displacements caused by pre- and post-saccadic eye drifts were removed and padded, to avoid temporal modulations from other types of eye movements. We then estimated *Q* directly in the frequency domain, as:

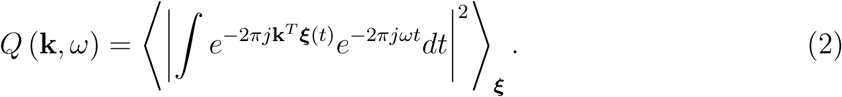

In Fig. 1*f*, the average redistributions across *N* =18 observers obtained over saccades with approximate amplitudes of 6*^◦^* and 1*^◦^* are plotted in two dimensions after averaging across spatial frequency, *Q*(*k, ω*) =*< Q*(**k***, ω*) *>_|_***_k_***_|_*_=_*_k_*.

To examine the power delivered by saccades within the range of human temporal sensitivity, we weighted *Q*(*k, ω*) by the temporal sensitivity function *G* and integrated across temporal frequencies.

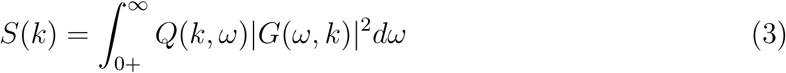

where *G* was taken from classical psychophysical measurements obtained under retinal stabilization ^43^, a condition that eliminates possible confounds from eye movements:

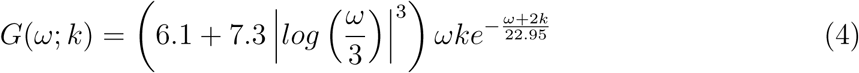

To provide an intuition of the expected consequences of power redistributions from saccades, we modeled the average response of neuronal populations tuned to the orientation and frequencies of the Probe and the Reference:

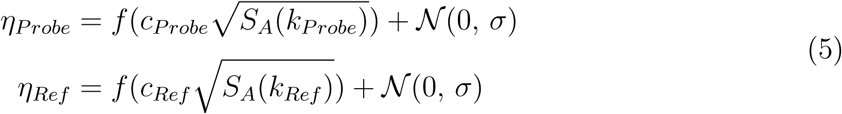

where *c* is the contrast of the grating; *S_A_*(*k*) represents the strength of the visual input signals resulting from a saccade of amplitude *A* during viewing of a stimulus at spatial frequency *k*; *f* is a standard logistic function; and N Gaussian noise with *σ* = 0.1. *S_A_*(*k*) was estimated for each individual observer, based on their recorded saccades. In each trial, the model compared the two responses and reported as most visible the grating that elicited the highest activity. The resulting psychometric functions evaluated over 30 contrast levels and 100 trials per level are reported in Fig. 2d.

## Supporting information

SuppFigs

## Acknowledgments

This work was supported by Reality Labs and grants R01 EY18363, EY07977, and P30 EY001319 from the National Institutes of Health. We thank Scott Murdison, Martina Poletti, and the members of the Active Perception Laboratory at the University of Rochester for helpful discussions.

## Author contributions

JDV and MR conceived the study. YHL implemented the experiments and collected the data. YHL and MAC analyzed the data. All authors contributed to the interpretation of the results. YHL, JDV and MR wrote the article. MR supervised the project.

## References

1. Vliegen, J., Van Grootel, T. J. & Van Opstal, A. J. Dynamic sound localization during rapid eye-head gaze shifts. J. Neurosci. 24, 9291–9302 (2004).

2. Kondo, H. M., Pressnitzer, D., Toshima, I. & Kashino, M. Effects of self-motion on auditory scene analysis. Proc. Natl. Acad. Sci. USA 109, 6775–6780 (2012).

3. Szwed, M., Bagdasarian, K. & Ahissar, E. Encoding of vibrissal active touch. Neuron 40, 621–630 (2003).

4. Kleinfeld, D., Ahissar, E. & Diamond, M. E. Active sensation: Insights from the rodent vibrissa sensorimotor system. Curr. Opin. Neurobiol. 16, 435–444 (2006).

5. Wachowiak, M. Active sensing in olfaction. In Menini, A. (ed.) The Neuroscience of Olfaction (CRC Press/Taylor & Francis, 2010).

6. Gibson, E. J., Gibson, J. J., Smith, O. W. & Flock, H. Motion parallax as a determinant of perceived depth. J. Exp. Psychol. 58, 40–51 (1959).

7. Rogers, B. & Graham, M. Motion parallax as an independent cue for depth perception. Perception 8, 125–134 (1979).

8. Howard, I. P. Perceiving in Depth, Vol. 3. Other Mechanisms of Depth Perception (Oxford University Press, 2012).

9. Averill, H. I. & Weymouth, F. W. Visual perception and the retinal mosaic. II. The influence of eye movements on the displacement threshold. J. Comp. Psychol. 5, 147– 176 (1925).

10. Marshall, W. H. & Talbot, S. A. Recent evidence for neural mechanisms in vision leading to a general theory of sensory acuity. In Kluver, H. (ed.) Biological Symposia— Visual Mechanisms, vol. 7, 117–164 (Cattel, Lancaster, PA, 1942).

11. Arend, L. E. Spatial differential and integral operations in human vision: Implications of stabilized retinal image fading. Psychol. Rev. 80, 374–395 (1973).

12. Ahissar, E. & Arieli, A. Figuring space by time. Neuron 32, 185–201 (2001).

13. Greschner, M., Bongard, M., Rujan, P. & Ammermuller, J. Retinal ganglion cell synchronization by fixational eye movements improves feature estimation. Nat. Neurosci. 5, 341–347 (2002).

14. Rucci, M., Ahissar, E. & Burr, D. Temporal coding of visual space. Trends Cogn. Sci. 22, 883–895 (2018).

15. Robson, J. G. Spatial and temporal contrast-sensitivity functions of the visual system. J. Opt. Soc. Am. 56, 1141–1142 (1966).

16. Purpura, K., Tranchina, D., Kaplan, E. & Shapley, R. M. Light adaptation in the primate retina: Analysis of changes in gain and dynamics of monkey retinal ganglion cells. Vis. Neurosci. 4, 75–93 (1990).

17. Kaplan, E. & Benardete, E. The dynamics of primate retinal ganglion cells. Prog. Brain. Res. 134, 17–34 (2001).

18. Derrington, A. M. & Lennie, P. Spatial and temporal contrast sensitivities of neurones in lateral geniculate nucleus of macaque. J. Physiol. 357, 219–240 (1984).

19. Wu, E. G. et al. Fixational eye movements enhance the precision of visual information transmitted by the primate retina. Nat. Commun. 15 (2024).

20. Ratliff, F. & Riggs, L. A. Involuntary motions of the eye during monocular fixation. J. Exp. Psychol. 40, 687–701 (1950).

21. Barlow, H. B. Eye movements during fixation. J. Physiol. 116, 290–306 (1952).

22. Kowler, E. Eye movements: The past 25 years. Vision Res. 51, 1457–1483 (2011).

23. Kuang, X., Poletti, M., Victor, J. D. & Rucci, M. Temporal encoding of spatial information during active visual fixation. Curr. Biol. 20, 510–514 (2012).

24. Rucci, M., Iovin, R., Poletti, M. & Santini, F. Miniature eye movements enhance fine spatial detail. Nature 447, 852–855 (2007).

25. Ratnam, K., Domdei, N., Harmening, W. M. & Roorda, A. Benefits of retinal image motion at the limits of spatial vision. J. Vis. 17, 30 (2017).

26. Intoy, J. & Rucci, M. Finely tuned eye movements enhance visual acuity. Nat. Commun. 11, 1–11 (2020).

27. Clark, A. M., Intoy, J., Rucci, M. & Poletti, M. Eye drift during fixation predicts visual acuity. Proc. Natl. Acad. Sci. USA 119, 1–10 (2022).

28. Intoy, J. et al. Consequences of eye movements for spatial selectivity. Curr. Biol. 34, 3265–3272 (2024).

29. Purpura, K. P., Kalik, S. F. & Schiff, N. D. Analysis of perisaccadic field potentials in the occipitotemporal pathway during active vision. J. Neurophysiol. 90, 3455–3478 (2003).

30. Kagan, I., Gur, M. & Snodderly, D. M. Saccades and drifts differentially modulate neuronal activity in V1: Effects of retinal image motion, position, and extraretinal influences. J. Vis. 8, 1–25 (2008).

31. Ibbotson, M. R., Crowder, N. A., Cloherty, S. L., Price, N. S. C. & Mustari, M. J. Saccadic modulation of neural responses: Possible roles in saccadic suppression, enhancement, and time compression. J. Neurosci. 28, 10952–10960 (2008).

32. Bremmer, F., Kubischik, M., Hoffmann, K. P. & Krekelberg, B. Neural dynamics of saccadic suppression. J. Neurosci. 29, 12374–12383 (2009).

33. Ruiz, O. & Paradiso, M. A. Macaque V1 representations in natural and reduced visual contexts: Spatial and temporal properties and influence of saccadic eye movements. J. Neurophysiol. 108, 324–333 (2012).

34. Bahill, A. T., Clark, M. R. & Stark, L. The main sequence, a tool for studying human eye movements. Math. Biosci. 24, 191–204 (1975).

35. Gibaldi, A. & Sabatini, S. P. The saccade main sequence revised: A fast and repeatable tool for oculomotor analysis. Behav. Res. Methods 53, 167–187 (2021).

36. Mostofi, N. et al. Spatiotemporal content of saccade transients. Curr. Biol. 30, 3999– 4008 (2020).

37. Mostofi, N., Boi, M. & Rucci, M. Are the visual transients from microsaccades helpful? Measuring the influences of small saccades on contrast sensitivity. Vision Res. 118, 60–69 (2016).

38. Navon, D. Forest before trees: The precedence of global features in visual perception. Cogn. Psychol. 9, 353–383 (1977).

39. Watt, R. J. Scanning from coarse to fine spatial scales in the human visual system after the onset of a stimulus. J. Opt. Soc. Am. A 4, 2006–2021 (1987).

40. Schyns, P. G. & Oliva, A. From blobs to boundary edges: Evidence for time- and spatial-scale-dependent scene recognition. Psychol. Sci. 5, 195–200 (1994).

41. Neri, P. Coarse to fine dynamics of monocular and binocular processing in human pattern vision. Proc. Natl. Acad. Sci. USA 108, 10726–10731 (2011).

42. Boi, M., Poletti, M., Victor, J. D. & Rucci, M. Consequences of the oculomotor cycle for the dynamics of perception. Curr. Biol. 27, 1–10 (2017).

43. Kelly, D. H. Motion and vision. II. Stabilized spatio-temporal threshold surface. J. Opt. Soc. Am. 69, 1340–1349 (1979).

44. Lin, Y., Intoy, J., Clark, A. M., Rucci, M. & Victor, J. D. Cognitive influences on fixational eye movements. Curr. Biol. 33, 1606–1612 (2023).

45. Carandini, M. & Heeger, D. J. Normalization as a canonical neural computation. Nat. Rev. Neurosci. 13, 51–62 (2012).

46. O’Regan, J. K. & Nöe, A. A sensorimotor account of vision and visual consciousness. Behav. Brain Sci. 24, 939–973 (2001).

47. Watanabe, A., Mori, T., Nagata, S. & Hiwatashi, K. Spatial sine-wave responses of the human visual system. Vision Res. 8, 1245–1263 (1968).

48. Georgeson, M. A. & Sullivan, G. D. Contrast constancy: Deblurring in human vision by spatial frequency channels. J. Physiol. 252, 627–656 (1975).

49. Knöll, J., Binda, P., Morrone, M. C. & Bremmer, F. Spatiotemporal profile of perisaccadic contrast sensitivity. J. Vis. 11, 15 (2011).

50. Binda, P. & Morrone, M. C. Vision during saccadic eye movements. Annu. Rev. Vis. Sci. 4, 193–213 (2018).

51. Intoy, J., Mostofi, N. & Rucci, M. Fast and nonuniform dynamics of perisaccadic vision in the central fovea. Proc. Natl. Acad. Sci. USA 118, e2101259118 (2021).

52. Gellman, R. S., Carl, J. R. & Miles, F. A. Short latency ocular-following responses in man. Vis. Neurosci. 5, 107–122 (1990).

53. Busettini, C., Miles, F. A. & Krauzlis, R. J. Short-latency disparity vergence responses and their dependence on a prior saccadic eye movement. J. Neurophysiol. 75, 1392– 1410 (1996).

54. Kawano, K. & Miles, F. A. Short-latency ocular following responses of monkey. II. Dependence on a prior saccadic eye movement. J. Neurophysiol. 56, 1355–1380 (1986).

55. Deubel, H. The time course of presaccadic attention shifts. Psychol. Res. 72, 630–640 (2008).

56. Li, H. H., Pan, J. & Carrasco, M. Presaccadic attention improves or impairs performance by enhancing sensitivity to higher spatial frequencies. Sci. Rep. 9 (2019).

57. Rolfs, M. & Carrasco, M. Rapid simultaneous enhancement of visual sensitivity and perceived contrast during saccade preparation. J. Neurosci. 32, 13744–13752 (2012).

58. Lisberger, S. G. Postsaccadic enhancement of initiation of smooth pursuit eye movements in monkeys. J. Neurophysiol. 79, 1918–1930 (1998).

59. Ramcharan, E. J., Gnadt, J. W. & Sherman, S. M. The effects of saccadic eye movements on the activity of geniculate relay neurons in the monkey. Vis. Neurosci. 18, 253–258 (2001).

60. Reppas, J. B., Usrey, W. M. & Reid, R. C. Saccadic eye movements modulate visual responses in the lateral geniculate nucleus. Neuron 35, 961–974 (2002).

61. Ibbotson, M. & Krekelberg, B. Visual perception and saccadic eye movements. Curr. Opin. Neurobiol. 21, 553–558 (2011).

62. Rolfs, M. & Schweitzer, R. Coupling perception to action through incidental sensory consequences of motor behavior. Nat. Rev. Psychol. 1, 112–123 (2022).

63. Miles, F., Kawano, K. & Optican, L. Short-latency ocular following responses of monkey. I. Dependence on temporospatial properties of visual input. J. Neurophysiol. 56, 1321–1354 (1986).

64. Watson, T. L. & Krekelberg, B. The relationship between saccadic suppression and perceptual stability. Curr. Biol. 19, 1040–1043 (2009).

65. Schweitzer, R. & Rolfs, M. Intra-saccadic motion streaks as cues to linking object locations across saccades. J. Vis. 20, 1–24 (2020).

66. Zimmermann, E. & Lappe, M. Eye position effects in oculomotor plasticity and visual localization. J. Neurosci. 31, 7341–7348 (2011).

67. Volkmann, F. C. Vision during voluntary saccadic eye movements. J. Opt. Soc. Am. 52, 571–578 (1962).

68. Campbell, F. W. & Wurtz, R. H. Saccadic omission: Why we do not see a grey-out during a saccadic eye movement. Vision Res. 10, 1297–1303 (1978).

69. Wurtz, R. H. Neuronal mechanisms of visual stability. Vision Res. 48, 2070–2089 (2008).

70. Volkmann, F. C. Human visual suppression. Vision Res. 26, 1401–1416 (1986).

71. Burr, D. C., Morrone, M. C. & Ross, J. Selective suppression of the magnocellular visual pathway during saccadic eye movements. Nature 371, 511–513 (1994).

72. Wurtz, R. H. Corollary discharge contributions to perceptual continuity across saccades. Annu. Rev. Vis. Sci. 4, 215–237 (2018).

73. Diamond, M. R., Ross, J. & Morrone, M. C. Extraretinal control of saccadic suppression. J. Neurosci. 20, 3449–3455 (2000).

74. MacKay, D. M. Elevation of visual threshold by displacement of retinal image. Nature 225, 90–92 (1970).

75. Castet, E., Jeanjean, S. & Masson, G. S. ‘Saccadic suppression’ – no need for an active extra-retinal mechanism. Trends Neurosci. 24, 316–317 (2001).

76. Idrees, S., Baumann, M. P., Franke, F., Munch, T. A. & Hafed, Z. M. Perceptual saccadic suppression starts in the retina. Nat. Commun. 11, 1–19 (2020).

77. Volkmann, F. C., Riggs, L. A., White, K. D. & Moore, R. K. Contrast sensitivity during saccadic eye movements. Vision Res. 18, 1193–1199 (1978).

78. Berman, R. A., Cavanaugh, J., McAlonan, K. & Wurtz, R. H. A circuit for saccadic suppression in the primate brain. J. Neurophysiol. 117, 1720–1735 (2017).

79. Burr, D. C. & Ross, J. Contrast sensitivity at high velocities. Vision Res. 22, 479–484 (1982).

80. Stevenson, S. B., Volkmann, F. C., Kelly, J. P. & Riggs, L. A. Dependence of visual suppression on the amplitudes of saccades and blinks. Vision Res. 26, 1815–1824 (1986).

81. Rothkegel, L. O. M., Schutt, H. H., Trukenbrod, H. A., Wichmann, F. A. & Engbert, R. Searchers adjust their eye-movement dynamics to target characteristics in natural scenes. Sci. Rep. 9, 1635 (2019).

82. Cajar, A., Engbert, R. & Laubrock, J. Spatial frequency processing in the central and peripheral visual field during scene viewing. Vision Res. 127, 186–197 (2016).

83. de Brouwer, A. J., Flanagan, J. R. & Spering, M. Functional use of eye movements for an acting system. Trends Cogn. Sci. 25, 252–263 (2021).

84. Antoniades, C. A. & Spering, M. Eye movements in Parkinson’s disease: From neurophysiological mechanisms to diagnostic tools. Trends Neurosci. 47, 71–83 (2024).

85. McDowell, J. E., Clementz, B. A. & Wixted, J. T. Timing and amplitude of saccades during predictive saccadic tracking in schizophrenia. Psychophysiology 33, 93–101 (1996).

86. Rommelse, N. N. J., Van der Stigchel, S. & Sergeant, J. A. A review on eye movement studies in childhood and adolescent psychiatry. Brain Cogn. 68, 391–414 (2008).

87. Obyedkov, I. et al. Saccadic eye movements in different dimensions of schizophrenia and in clinical high-risk state for psychosis. BMC Psychiatry 19, 1–10 (2019).

88. Lencer, R. et al. Saccadic suppression in schizophrenia. Sci. Rep. 11, 13133 (2021).

89. Butler, P. D., Silverstein, S. M. & Dakin, S. C. Visual perception and its impairment in schizophrenia. Biol. Psychiatry. 64, 40–47 (2008).

90. Shoham, N. et al. Investigating the association between schizophrenia and distance visual acuity: Mendelian randomisation study. BJPsych. Open. 9, e33 (2023).

91. Schmitt, L. M., Cook, E. H., Sweeney, J. A. & Mosconi, M. W. Saccadic eye movement abnormalities in autism spectrum disorder indicate dysfunctions in cerebellum and brainstem. Mol. Autism 5, 47 (2014).

92. Kéïta, L., Guy, J., Berthiaume, C., Mottron, L. & Bertone, A. An early origin for detailed perception in autism spectrum disorder: Biased sensitivity for high-spatial frequency information. Sci. Rep. 4, 5475 (2014).

93. Santini, F., Redner, G., Iovin, R. & Rucci, M. EyeRIS: A general-purpose system for eye movement contingent display control. Behav. Res. Methods 39, 350–364 (2007).

94. Wu, R.-J. et al. High-resolution eye-tracking via digital imaging of Purkinje reflections. J. Vis. 23, 4 (2023).

95. Poletti, M. & Rucci, M. A compact field guide to the study of microsaccades: Challenges and functions. Vision Res. 118, 83–97 (2016).

96. Wichmann, F. A. & Hill, N. J. The psychometric function: I. Fitting, sampling, and goodness of fit. Percept. Psychophys. 63, 1293–1313 (2001).

97. Hall, J. L. Hybrid adaptive procedure for estimation of psychometric functions. J. Acoust. Soc. Am. 69, 1763–1769 (1981).

