## Supplementary material for "What one sees depends on how far the eye has moved": SuppFigs

### Supplementary figures

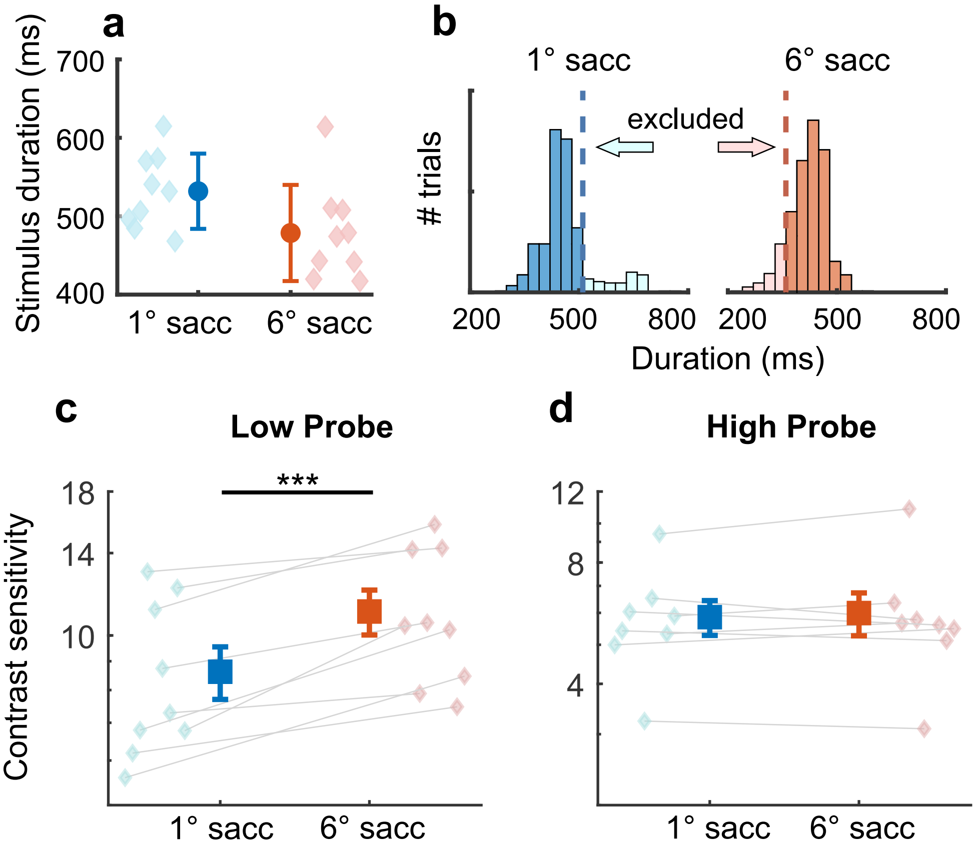


Figure S1: Control for pre-saccadic stimulus exposure. **(A)** Mean durations of the intervals of stimulus exposure before performing saccades of either 1° or 6° amplitude. On average, exposure was 53 ms longer with the smaller saccade. Diamonds mark the mean exposure duration for each subject; circles represent averages across subjects. **(B)** Histograms of pre-saccadic stimulus exposure durations for a representative subject. The light-shaded regions mark the trials discarded to compensate for differences in saccade reaction times. **(C-D)** Sensitivity in the two conditions changes little after equalizing pre-saccadic exposure ($p < 0.001$ for Low-Probe; $p = 0.59$ for High-Probe).

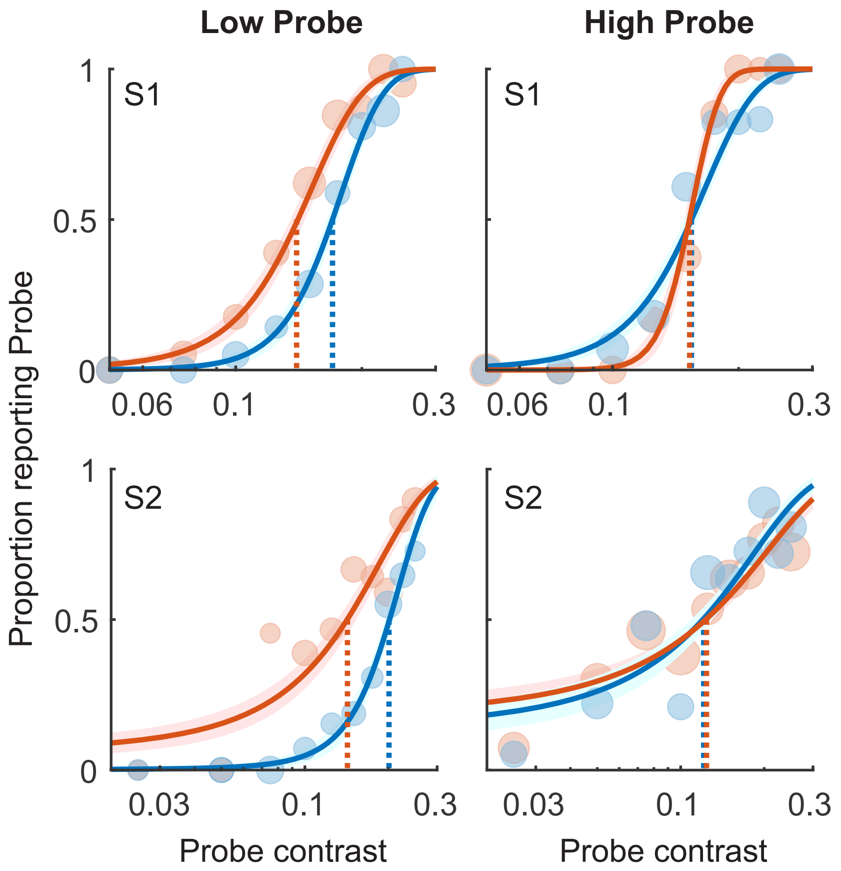


Figure S2: Psychometric functions measured during exposure to simulated saccades. In this experiment, subjects maintained fixation for the entire duration of the trial. The stimulus moved on the display following previously recorded saccade traces to replicate the visual input signals normally delivered by saccades. Each row shows data from one subject.

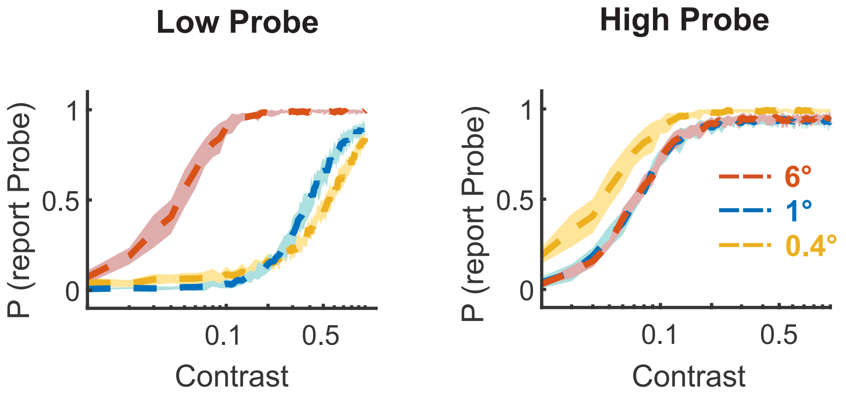


Figure S3: Psychometric functions predicted by the ideal-observer model when also considering the 0.4° saccade. A perceptual enhancement for the 0.4° saccade is expected in the High-Probe but not in the Low-Probe condition.
